# Meiotic drive reduces egg-to-adult viability in stalk-eyed flies

**DOI:** 10.1101/690321

**Authors:** Sam Ronan Finnegan, Nathan Joseph White, Dixon Koh, M. Florencia Camus, Kevin Fowler, Andrew Pomiankowski

**Author notes:** **Corresponding author:** Andrew Pomiankowski.

## Abstract

SR meiotic drive is a selfish genetic element located on the X chromosome in a number of species that causes dysfunction of Y-bearing sperm. SR is transmitted to up to 100% of offspring, causing extreme sex ratio bias. SR in several species is found in a stable polymorphism at a moderate frequency, suggesting there must be strong frequency-dependent selection resisting its spread. We investigate the effect of SR on female and male egg-to-adult viability in the Malaysian stalk-eyed fly, *Teleopsis dalmanni*. SR meiotic drive in this species is old, and appears to be broadly stable at a moderate (~20%) frequency. We use large-scale controlled crosses to estimate the strength of selection acting against SR in female and male carriers. We find that SR reduces the egg-to-adult viability of both sexes. In females, homozygous females experience greater reduction in viability (*s*_*f*_ = 0.242) and the deleterious effects of SR are additive (ℎ = 0.511). The male deficit in viability (*s*_*m*_ = 0.214) is not different from that in homozygous females. The evidence does not support the expectation that deleterious side-effects of SR are recessive or sex-limited. We discuss how these reductions in egg-to-adult survival, as well as other forms of selection acting on SR, act to maintain SR polymorphism in this species.

## Introduction

Meiotic drivers are selfish genetic elements that subvert the standard mechanisms of gametogenesis to promote their own transmission (Lindholm et al. 2016). During meiosis, a driver disables or prevents the maturation of gametes that contain the non-driving element (Burt and Trivers 2006; Lindholm et al. 2016). In extreme cases, drive can reach 100% transmission to the next generation. In male heterogametic species, drivers are most frequently found on the X-chromosome (Hurst and Pomiankowski 1991), commonly known as ‘*Sex-Ratio*’ or SR (Hurst & Werren 2001). These drivers target developing sperm carrying the Y chromosome, causing their dysfunction, which results in strongly female biased broods.

SR is predicted to spread rapidly due to its transmission advantage. When homozygous female fitness is not greatly reduced, SR could potentially spread to fixation and cause population collapse and extinction through massive sex ratio imbalance (Hamilton 1967, Hatcher et al. 1999). Empirical evidence for this is limited to laboratory environments where drive causes extinction in small populations (Lyttle 1977, Price et al. 2010, Galizi et al. 2014) and a single putative example under natural conditions (Pinzone & Dyer 2013). More typically, studies in wild populations find that drive exists as a low-frequency polymorphism (Pinzone & Dyer 2013; Manser et al. 2011; Price et al. 2014; Verspoor et al. 2018), with persistence that can span over a million years (Silver 1993; Kovacevic & Schaeffer 2000; Paczolt et al. 2017). In order for SR to persist as a polymorphism, there must be frequency-dependent selection, allowing spread when rare but retarding further increases in frequency as drive becomes more common. The selective counter forces that fulfil this requirement may act in males or females but in general they are not well understood. We discuss potential causes of selection first in males and then females in the following sections.

Selection on male viability may be associated with the drive chromosome. It is likely to operate in a frequency-independent manner and so not have a stabilizing effect on the frequency of drive (Edwards 1961; Carvalho and Vaz 1999). But it has been suggested that there will be negative frequency-dependent selection on male fertility (Jaenike 1996). This has intuitive appeal because the spread of SR causes the population sex ratio to become increasingly female biased. In such a population, the average male mating rate will increase. If SR male fertility increases at a lower rate than non-drive (ST) male fertility when males mate multiply (for instance because SR males are sperm limited), then a polymorphism could be stabilised (Jaenike 1996). Decreased male fertility under multiple mating is a general feature observed in many drive systems (Beckenbach, 1978; Jaenike, 1996; Atlan et al. 2004). However, for this effect alone to prevent SR fixation, SR male fertility must fall to less than half that of ST males as the mating rate increases (Jaenike 1996), a condition not met in a number of species that nonetheless are found with stable SR polymorphism (Carvalho and Vaz 1999). A related suggestion is that SR males may be out-competed at higher mating rates, motivated by some evidence that SR males are poor sperm competitors (Wu 1983a; Wilkinson and Fry 2001; Price and Wedell 2008). However, the strength of sperm competition weakens as SR spreads, as this reduces the number of competitor males in the population, which seems unlikely to exert a stabilizing effect on SR frequency. SR males may do poorly in other forms of male-male competition if SR is generally associated with poor performance. Such effects are likely to decrease as drive spreads and males become rare, again making it unlikely that this form of selection will stabilize drive. Models that combine the effects of decreased male fertility and reduced sperm competitive ability on SR frequency dynamics find they can lead to a stable polymorphism (Taylor and Jaenike 2002). But this equilibrium can be destabilised by perturbations in either the population sex ratio or the frequency of SR. In particular, given a meta-population of small demes, slight fluctuations in SR frequency are likely to cause drive to spread to fixation, resulting in population extinction (Taylor and Jaenike 2003).

Suppressors are another selective force operating in males that limits the spread of drive alleles. Most obviously, selection favours the evolution of suppression on chromosomes targeted by drivers for dysfunction. In an SR system with complete drive, if resistance is linked to the Y-chromosome, it restores transmission to Mendelian levels, while non-resistant Y-chromosomes are not transmitted at all (Thomson and Feldman 1975). Y-linked suppressors are therefore expected to spread quickly even if they have deleterious side effects (Wu 1983b). Unlinked suppressors will also be favoured because drive in males causes gamete loss and is often associated with dysfunction amongst the surviving, drive-carrying sperm. Reduced sperm number is likely to reduce organismal fertility. Additionally, as SR spreads it causes the population sex ratio to become female-biased, providing a further advantage to suppressors as they increase the production of male offspring, which have higher reproductive value than female offspring (Fisher 1930; Carvalho et al. 1998). The spread of suppressors reduces the advantage of drive and could lead to its loss. But both types of suppressors are under negative frequency-dependent selection, because a lower frequency of drive reduces selection in their favour. Under some circumstances this could lead to a stable polymorphism at the drive locus. Y-linked and autosomal suppressors of SR drive have been detected in a number of species including *D. simulans*, *D.affinis, D. subobscura, D. quinara, D. mediopunctata* and *Aedes aegypti* (Jaenike 2001). The evolution of suppressors can be remarkably rapid. For example, in the Paris SR system of *D. simulans*, the increase of SR from less than 10% to more than 60% in a mere five years has been matched by a similar increase in suppressor frequency over the same time period (Bastide at al. 2013). While suppressors are common, they are not universal and have not been detected against SR drive in *D. pseudoobscura, D. recens* and *D. neotestacea* (Jaenike 2001). In these systems, other factors are therefore necessary to explain extant SR polymorphism.

Alternatively, SR may be prevented from reaching fixation if female carriers have reduced fitness (Curtsinger and Feldman 1980). As male X-linked drive causes defects in spermatogenesis, there is no obvious mechanistic carry-over to female oogenesis. Likewise, examples of meiotic drive in female gametogenesis, which affect the biased segregation of chromosomes into the egg or polar bodies, show no carry-over to segregation bias in male gamete production (Burt and Trivers 2006). For selection to act against female carriers, the drive locus must either have direct pleiotropic fitness effects or be in linkage with alleles that impact fitness. Linkage is a plausible explanatory factor given that drive systems are often located in genomic regions with low recombination rates, such as in inversions (Beckenbach 1996; Silver 1993; Dyer et al. 2007; Reinhardt et al. 2014). If the inversion is at low frequency, it will rarely be homozygous and the recombination rate among SR chromosomes will be low. Inversions also severely limit the exchange of genes with the homologous region on the standard chromosome (as this requires a double cross-over within the inverted region; Navarro et al. 1997; 1998). The consequence is that low frequency inversions will be subject to weak selection and suffer the accumulation of a greater mutation load (Dyer et al. 2007; Kirkpatrick 2010). Recessive viability and sterility effects are expected as they will not be evident in females until the frequency of drive is sufficient for the production of homozygotes. In contrast, hemizygosity in males means recessive and dominant effects are always expressed. This means that female-limited fitness effects are more likely to produce relevant frequency dependence that restricts fixation of drive. Severe reductions in female viability and fertility in SR homozygotes, along with SR heterozygotes, have been reported in several *Drosophila* species (Wallace 1948; Curtsinger and Feldman 1980; Dyer et al. 2007). But it is surprising how rarely viability effects of drive in either sex have been studied, compared to fertility effects in males (Price and Weddell 2008). These deleterious consequences are likely to build up and lead to a reduction in SR frequency through time (Dyer et al. 2007).

Large-scale chromosomal inversions are not a universal feature of SR, however. Inversions are not present in the Paris SR system in *D. simulans* (Jaenike 2001). Despite this, SR must be weakly deleterious in this species as it is rapidly declining in frequency in populations that have recently become completely suppressed (Bastide et al. 2011). The deleterious effects of the Paris SR chromosome must arise due to deleterious effects caused by the drive genes themselves or a tightly linked region. The genetically distinct Winters SR system in the same species also lacks association with an inversion (Kingan et al. 2010), It persists despite having been completely suppressed for thousands of years, suggesting it does not causes any pleiotropic fitness deficit (Kingan et al. 2010). These are the only well characterised examples of meiotic drive not being associated with inversions, so this feature may be a rarity.

Another aspect operating in females concerns behavioural resistance to the spread of SR. Laboratory experiments suggest that increased levels of polyandry can be selected as a defence mechanism against SR (Price et al. 2008). This benefit arises when drive male sperm are weak competitors against wildtype male sperm (Price and Wedell 2008). Recent modelling work shows that polyandry helps prevent invasion of SR, but cannot prevent fixation of drive alone (Holman et al. 2015). As drive spreads, additional matings have a lower probability of involving wildtype males, so the disadvantage to drive sperm declines. There needs to be positive frequency-dependent costs to achieve a stable polymorphism (Holman et al. 2015), for instance, when homozygous females have lower viability than heterozygotes. If a stable polymorphism can evolve, the frequency of drive should decline with the rate of female remating. There is evidence in favour of this idea in *D. neotestacea* which exhibits a stable cline in SR frequency that correlates negatively with the frequency of polyandry (Pinzone and Dyer, 2013), and a similar pattern has been reported in *D. pseudoobscura* (Price et al. 2014). Alternatively, females may simply avoid mating with SR males (Lande and Wilkinson 1999; Pomiankowski and Hurst 1999). In stalk-eyed flies, females prefer to mate with males with large eyespan (Wilkinson et al. 1998; Cotton et al. 2010), a trait that is reduced in SR males (Wilkinson et al. 1998; Johns et al. 2005; Cotton et al. 2014). Sexual selection may therefore be acting in this species to limit the spread of SR. However, this form of selection against drive is likely to be restricted to a sub-set of species with drive, as it requires the linkage of SR with a conspicuous trait subject to mate choice (Pomiankowski and Hurst 1999). Another potential example is the autosomal *t*-locus system in mice which is proposed to be detectable in mate choice through olfaction (Coopersmith and Lenington 1990) but this preference has not been confirmed (Sutter and Lindholm 2016). A counter example is in *D. pseudoobscura*, where females do not avoid mating with SR males, though there would be considerable benefit to doing so (Price et al. 2012).

In this study, we determine the effect of SR meiotic drive on viability in the Malaysian stalk-eyed fly, *Teleopsis dalmanni*. Our objective was to assess whether there is a SR-linked deleterious mutation load leading to higher developmental mortality before adult eclosion. Populations of this species carry SR at a moderate level of ~20% but with considerable variation among populations (Presgraves et al. 1997; Wilkinson et al. 2003; Paczolt et al. 2017). SR resides within a large paracentric inversion (or inversions) that covers most of the X chromosome (Johns et al. 2005). There is no recombination between SR and ST haplotypes (Paczolt et al. 2017) and the lower frequency of SR in the wild means SR and ST homozygous recombination events are relatively rare (at 20%, the recombination rate of SR is a quarter that of ST). SR is absent from a cryptic species of *T. dalmanni* estimated to have diverged ~1 Mya. X-linked meiotic drive is also present in the more distantly related species *T. whitei*, which diverged on order 2-3.5 Mya (Swallow et al. 2005; Paczolt et al. 2017). But to what extent the mechanism or genetic basis is conserved remains to be established.

The ancient origin of the X^SR^ chromosome and limited recombination across the X^SR^ chromosome are predicted to have led to the accumulation of deleterious alleles. The main evidence for this is the reduced eyespan of SR males (Wilkinson et al. 1998; Cotton et al. 2014). Male eyespan is an exaggerated, highly condition-dependent trait used in female mate choice (Wilkinson et al. 1998; Cotton et al. 2004), as well as signalling between males (Panhuis and Wilkinson 1999; Cotton et al. 2009), which reflects male genetic and phenotypic quality (David et al. 2000; Cotton et al. 2004; Howie et al. 2019). However, in a series of experiments Wilkinson et al. (2006) found little direct evidence that the SR reduces fitness components. Although larval viability was not direct assessed, progeny production showed no difference between SR and ST homozygous females (Wilkinson et al. 2006). Another study compared offspring genotypes of heterozygous females mated to ST males, and reported little deviation from 1:1 among SR:ST male offspring (Johns et al. 2005). Adult survival did not vary with genotype in either males or females (Wilkinson et al. 2006). There was no evidence for a deleterious effect of X^SR^ on female fecundity, rather heterozygotes were more productive, suggesting overdominance (Wilkinson et al. 2006). However, sample size in these experiments was small, and fecundity/fertility results were based on progeny counts which are confounded by genotype effects on larval survival. The only significant detriment reported was in SR male fertility which was reduced when males were allowed to mate with large numbers of females (eight) for 24 hours (Wilkinson et al. 2006). However, a further experiment that measured male fertility through counts of fertile eggs (avoiding any confounding impact of larval survival), failed to show any difference between SR and ST male fertility (Meade et al. 2019).

To better understand these previous results, we were motivated to explicitly test for differences in larval survival. Our experimental design was similar to that used in early investigations of *Drosophila pseudoobscura* (Wallace 1948; Curtsinger and Feldman 1980). Controlled crosses were carried out to produce eggs with of all possible SR and ST male and female genotypes. These were reared together to ensure exposure to similar environmental variation. The sample size was large to maximize our power to detect genotypic survival differences. Offspring were genotyped at adult eclosion, yielding observed genotype ratios in order to estimate the selection coefficients operating against drive in both sexes. Our principal aims were to test whether the SR-drive chromosome causes viability loss during egg-to-adult development, and whether fitness effects are recessive or sex-limited.

## Methods

### Fly stocks and maintenance

A standard stock population was obtained from Ulu Gombak in Malaysia (3°19’N 101°45’E) in 2005 (by Sam Cotton and Andrew Pomiankowski). Stock flies are reared in high-density cage culture (cage size approx. 30 × 20 × 20cm) at 25°C on a 12:12 hour light:dark cycle, and fed puréed corn *ad libitum*. Fifteen minute artificial dawn and dusk phases are created by illumination from a single 60-W at the start and end of each light phase. Meiotic drive is absent from the standard stock population.

A meiotic drive stock was created using flies collected from the same location in 2012 (Cotton et al 2014). Meiotic drive is maintained in this stock by following a standard protocol (Presgraves et al. 1997; Meade et al. 2018). Females heterozygous for the drive chromosome are mated to males from the standard stock. It is expected that half their male offspring will inherit the drive chromosome. All male offspring are crossed to three females from the standard stock and the sex ratio of their progeny scored. Males that sire all-female broods of at least 15 individuals are considered to be carriers of meiotic drive. In the meiotic drive stock, drive strength is 100% percent, and no males are produced by X^SR^/Y males carrying the drive chromosome (Meade et al. 2018). Progeny from drive males are female heterozygotes for the drive chromosome. They are subsequently mated to standard males, and the process is repeated.

### Experimental crosses

To generate the five possible genotypes of both females (X^ST^/X^ST^, X^SR^/X^ST^, X^SR^/X^SR^) and males (X^ST^/Y, X^SR^/Y), two crosses were performed (Figure 1). In Cross A, drive males (X^SR^/Y) are mated to heterozygous females (X^SR^/X). This cross produces X^SR^/X^SR^ and X^SR^/X^ST^ female zygotes in equal proportions. In Cross B, standard males (X^ST^/Y) are mated to heterozygous females (X^SR^/X^ST^). This cross produces X^ST^/Y and X^SR^/Y male, and X^ST^/X^ST^ and X^SR^/X female zygotes in equal proportions. Experimental males were collected from the drive stock that were approximately 50:50 X^ST^/Y and X^SR^/Y males. They were crossed to standard stock females (X^ST^/X^ST^) and one larva per male was genotyped to define the paternal genotype. Experimental females heterozygous for drive (X^SR^/X^ST^) were collected from crosses between drive males and females from the standard stock.

**Figure 1.**
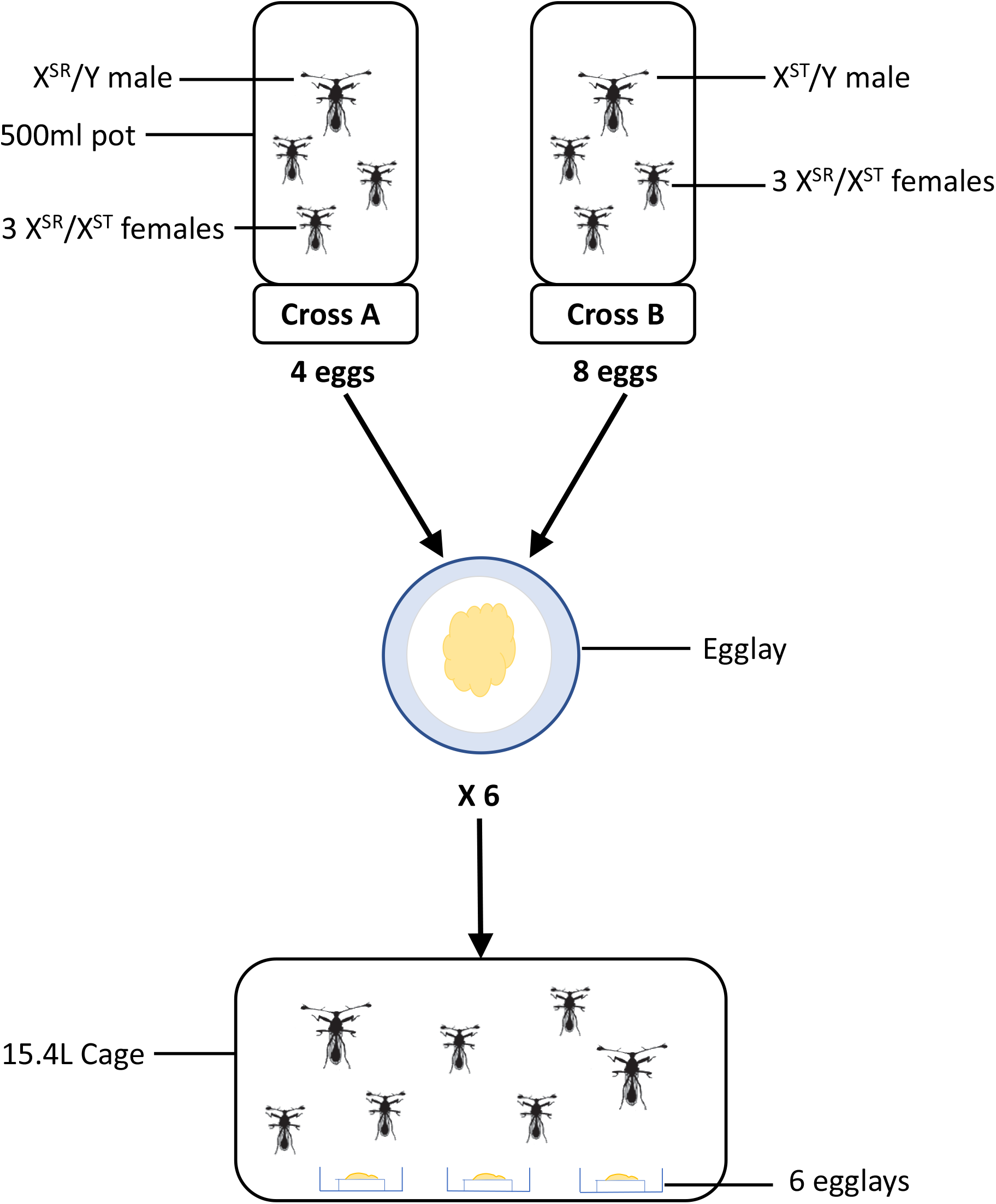
Experimental protocol. Individual males of known genotype were crossed with three heterozygous females in 500ml pots. Cross A produces no males and X^SR^/X^SR^ and X^SR^/X^ST^ females, in equal proportions. Cross B produces X^SR^/Y and X^ST^/Y males and X^ST^/X^ST^ and X^SR^/X^ST^ females, in equal proportions. 4 eggs from Cross A and 8 eggs from Cross B were added to each egglay – a petri dish containing a moistened cotton pad and food. At pupation, 6 egglays were placed into a population cage and their lids were removed so as to allow the adult flies to eclose.

Individual males were placed with three virgin females in 500ml pots. Females that died during the experiment were replaced, but males were not. 25 Cross A and 50 Cross B pots were set-up. The base of each pot was lined with moistened cotton wool covered with blue tissue paper to aid egg visualisation. The cotton bases were removed for egg collection and replaced three times per week. Fertilised eggs were identified under light microscopy as those that showed signs of development (e.g. segmental striations, development of mouthparts; Baker et al. 2001) and transferred to a 90mm petri dish containing a large cotton pad moistened with 15ml of water and 2.5ml of food. Three different food conditions were used that varied in their corn content: 25% corn, 50% corn, and 75% corn. In each mixture the remainder was made up with a sucrose solution (25% sucrose/water w/w). To ensure the sucrose solution had a similar viscosity to puréed corn, an indigestible bulking agent was added (methylcellulose, 3% w/w; Rogers et al. 2008). 4 eggs from Cross A and 8 eggs from Cross B were transferred to each petri dish. This gives the five possible genotypes (X^ST^/X^ST^, X^SR^/X^ST^, X^SR^/X^SR^, X^ST^/Y, X^SR^/Y) in an expected 1:2:1:1:1 ratio (Table 1). Prior to the end of development, six Petri dishes were placed inside a large cage and all eclosing adult flies were collected. The cage was used as a level of analysis of the relative egg-to-adult viability of different genotypes in the analysis that follows.

**Table 1:** Relative egg-to-adult viability. The five genotypes are drawn from crosses between heterozygous females and drive males (Cross A) or standard males (Cross B). The selection parameters, *s*_*f*_ and *s*_*m*_, measure drive egg-to-adult viability relative to wildtype females and males respectively. The dominance coefficient of drive is denoted *h*.

|  | Females |  |  | Males |  |
| --- | --- | --- | --- | --- | --- |
| | $X^{SR}/X^{SR}$ | $X^{SR}/X^{ST}$ | $X^{ST}/X^{ST}$ | $X^{SR}/Y$ | $X^{ST}/Y$ |
| **Cross A**<br>$X^{SR}/X^{ST} \times X^{SR}/Y$ | $1 - s_f$ | $1 - hs_f$ | | | |
| **Cross B**<br>$X^{SR}/X^{ST} \times X/Y$ | | $1 - hs_f$ | 1 | $1 - s_m$ | 1 |

### Genotyping

DNA was extracted by isopropanol precipitation in 96-well plates. Half a fly thorax was added to a well containing 4μl Proteinase K (10 mg.ml^−1^) and 100μl DIGSOL (25mM NaCl, 1mM EDTA, 10mM Tris–Cl pH 8.2), mechanically lysed, and incubated overnight at 55°C. The following day, 35μl of 4M ammonium acetate was added and plates were left on ice for 5 minutes before being centrifuged at 4500RPM at 4°C for 40 minutes. 80μl of supernatant was then aspirated into a new 96-well plate containing 80μl of isopropanol. The precipitate was discarded. Samples were then centrifuged again at 4500RPM and 4°C for 40 minutes to precipitate the DNA. The supernatant was then discarded, 100μl 70% ethanol was added, and samples were spun again at 4500RPM and 4°C for 20 minutes. The supernatant was once again discarded and plates were left to air-dry for 45 minutes at room temperature. Finally, 30μl of Low TE (1mM Tris-HCL pH8, 0.1mM EDTA) was added to elute the DNA. DNA was PCR-amplified in 96-well plates, with each well containing 1μl of dried DNA, 1μl of primer mix (consisting of the forward and reverse primers of *comp162710* at a concentration of 0.2μM) and 1μl of QIAGEN Multiplex PCR Mastermix (Qiagen). The length of amplified fragments was determined by gel electrophoresis. A 3% agarose gel was made using 3g of molecular grade agarose, 100ml of 0.5x TBE buffer (45mM Tris (pH 7.6), 45mM boric acid, 1mM EDTA), and 4μl ethidium bromide. PCR products were diluted with 3μl ultrapure water and 2μl of gel loading dye was added. 4μl of this mixture was loaded into each well and assessed for size against a ladder made from the PCR-amplified DNA of multiple heterozygous drive females. *comp162710* is an indel marker with small alleles (201bp) indicating the presence of the drive chromosome and large alleles (286bp) indicating the presence of the standard chromosome.

### Statistical analysis

We used two approaches to estimate the egg-to-adult viability costs of the X^SR^ chromosome. The first estimates the relative egg-to-adult viability cost of each genotype. The second estimates the strength of selection against drive in males and females, as well as the dominance coefficient.

### Egg-to-adult viability of each genotype

In the first analysis, the number of eclosed adult flies of each genotype were compared to the number expected at the level of the cage. Each cage contained six petri dishes with 12 eggs, producing a maximum of 72 flies. Genotyping effort varied across cages and sexes. The expected number of each genotype was determined with respect to the genotyping effort of the relevant sex for a particular cage. For example, if 75% of males in a given cage were genotyped, then the expected number of X^SR^ individuals is (0.75 × 72) / 6 = 8. We split the data by sex, then analysed the relationship between egg-to-adult viability and genotype using linear mixed-effect modelling with lme4 (Bates et al. 2015) in R (R Core Team, 2018). Genotype and food condition were modelled as fixed effects and cage ID and collection date as random effects. Significance of model terms was determined using the lmerTest R package (Kuznetsova et al. 2017). Food condition did not affect egg-to-adult viability, and so is not included in subsequent analyses. Mean viability measures were estimated using model terms.

### Estimating the strength of selection against drive

In the second analysis, we estimated the strength of selection against drive using Bayesian inference, separately for males and females. Cage survival frequencies for each genotype were pooled. The probability of drawing the male genotype distribution was calculated for values of the selection coefficient taken from a uniform prior distribution for *s*_*m*_ = 0 - 1, in 0.001 increments. We then used a binomial model to determine the likelihood of drawing the observed number of X^ST^/Y and X^SR^/Y males for each value of *s*_*m*_. As we used a uniform prior, the posterior probability simplifies to the likelihood. The 95% and 99% credible intervals were determined from the probability density. The probability of observing the distribution of the three female genotypes was estimated under a multinomial where the values of *s*_*f*_ and *h* (Table 1) were taken from a uniform prior distribution for every combination of values of *s*_*f*_ and *h* ranging from 0 - 1, in 0.001 intervals. The 95% and 99% credible intervals were determined in the same way as in males, and displayed as a two-dimensional contour. Note that the probability of drawing X^SR^/X^ST^ females was multiplied by two because the experimental design was expected ti generate twice as many heterozygote eggs compared to all of the other genotypes. To determine if *s*_*m*_ and *s*_*f*_ were of different strength, 1000 random samples each of *s*_*m*_ and *s*_*f*_ (taking *h* equal to its mode) were drawn from the posterior distributions with probability of drawing a value equal to its likelihood. A distribution of differences was obtained by subtracting the randomly drawn *s*_*f*_ values from the randomly drawn *s*_*m*_ values. A z-score was calculated to determine if this distribution is different from zero.

We also estimated the difference in the strength of selection between female genotypes. To compare egg-to-adult viability between wildtype (X^ST^/X^ST^) and heterozygous (X^SR^/X^ST^) females, the likelihood of observing the counts of these two genotypes was determined under a binomial as above, but shrinking *h* and *s*_*f*_ to a single term with a uniform prior. The process was repeated to compare drive heterozygotes (X^SR^/X^ST^) and homozygotes (X^SR^/X^SR^).

## Results

### Effect of food condition

Food condition had no overall effect on the egg-to-adult viability of males (F_2,72_ = 0.1085, P = 0.8973) or females (F_2,54_ = 0.1552, P = 0.9355), nor did it alter the genotype response (genotype-by-condition interaction, males F_2,79_ = 0.8026, P = 0.4518; females F_4,116_ = 0.2044, P = 0.9355). So, offspring counts were pooled across food conditions within sexes in the following analyses.

### Egg-to-adult viability of each genotype

We collected a total of 1065 males and 2500 females, of which 798 and 1272 were genotyped respectively. Male genotype had a significant effect on egg-to-adult viability, with X^SR^/Y males showing significantly reduced viability (F_1,81_ = 11.7296, P < 0.001). X^ST^/Y males had a mean viability of 0.5412, and X^SR^/Y males had a mean viability of 0.4036 (Figure 2). Genotype also had a significant effect on egg-to-adult viability in females (F_2,120_ = 4.7593, P = 0.0103). Mean viability was 0.6338 in X^ST^/X^ST^ females, 0.5537 in X^SR^/X^ST^ females, and 0.4695 in X^SR^/X^SR^ individuals. A Tukey’s post-hoc comparison test revealed that the viability of X^ST^/X^ST^ females is greater than X^SR^/X^SR^ females (P = 0.0109), while X^SR^/X^ST^ females have intermediate viability, but not different from either homozygote (X^SR^/X^ST^ – X^SR^/X^SR^ comparison: P = 0.2949; X^SR^/X^ST^ – X^ST^/X^ST^ comparison: P = 0.3293; Figure 3).

**Figure 2.**
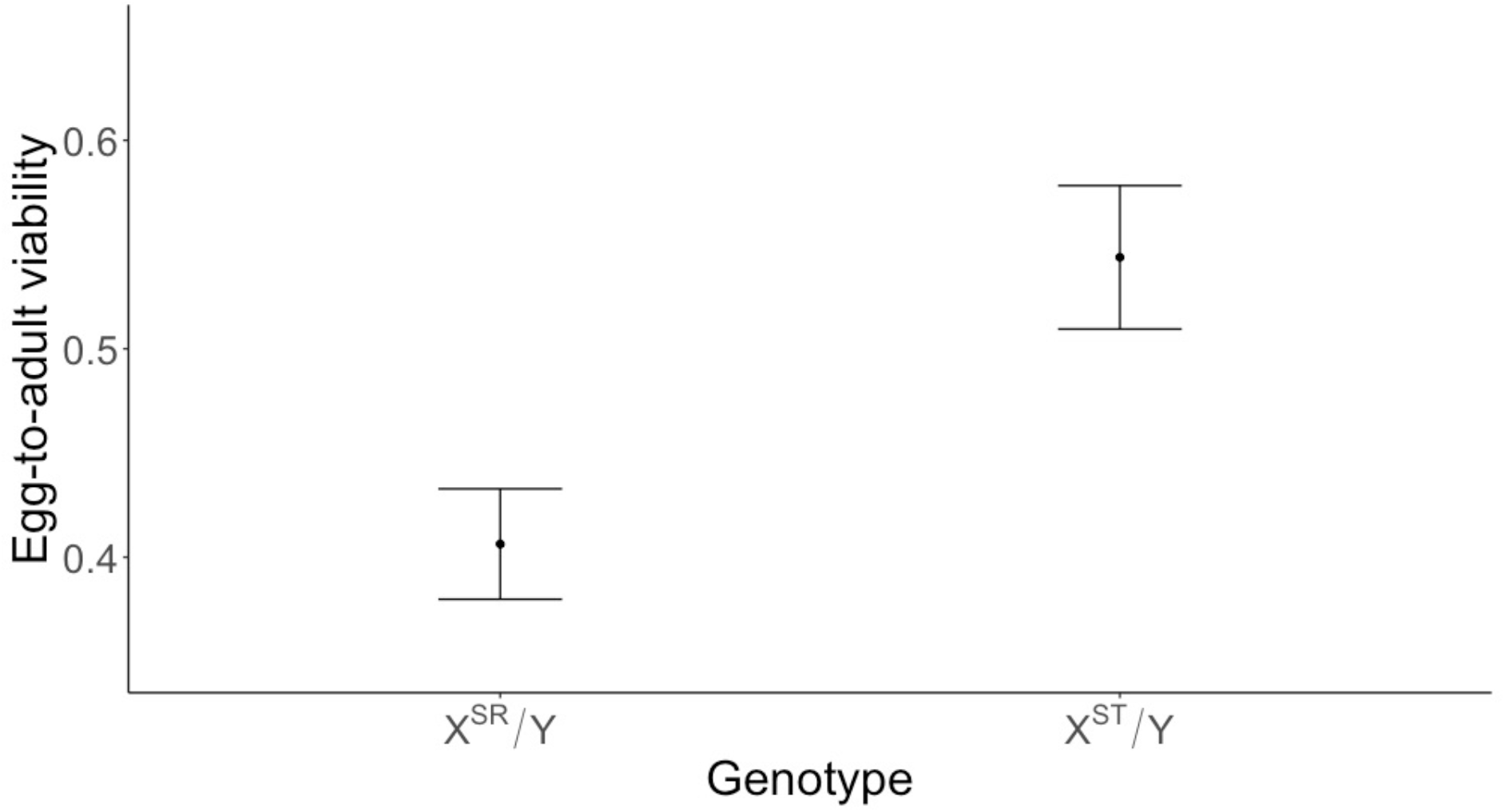
Male X^SR^/Y and X^ST^/Y mean ± standard error proportion egg-to-adult viability. Values are determined from the fraction of a given genotype observed in replicate cages.

**Figure 3.**
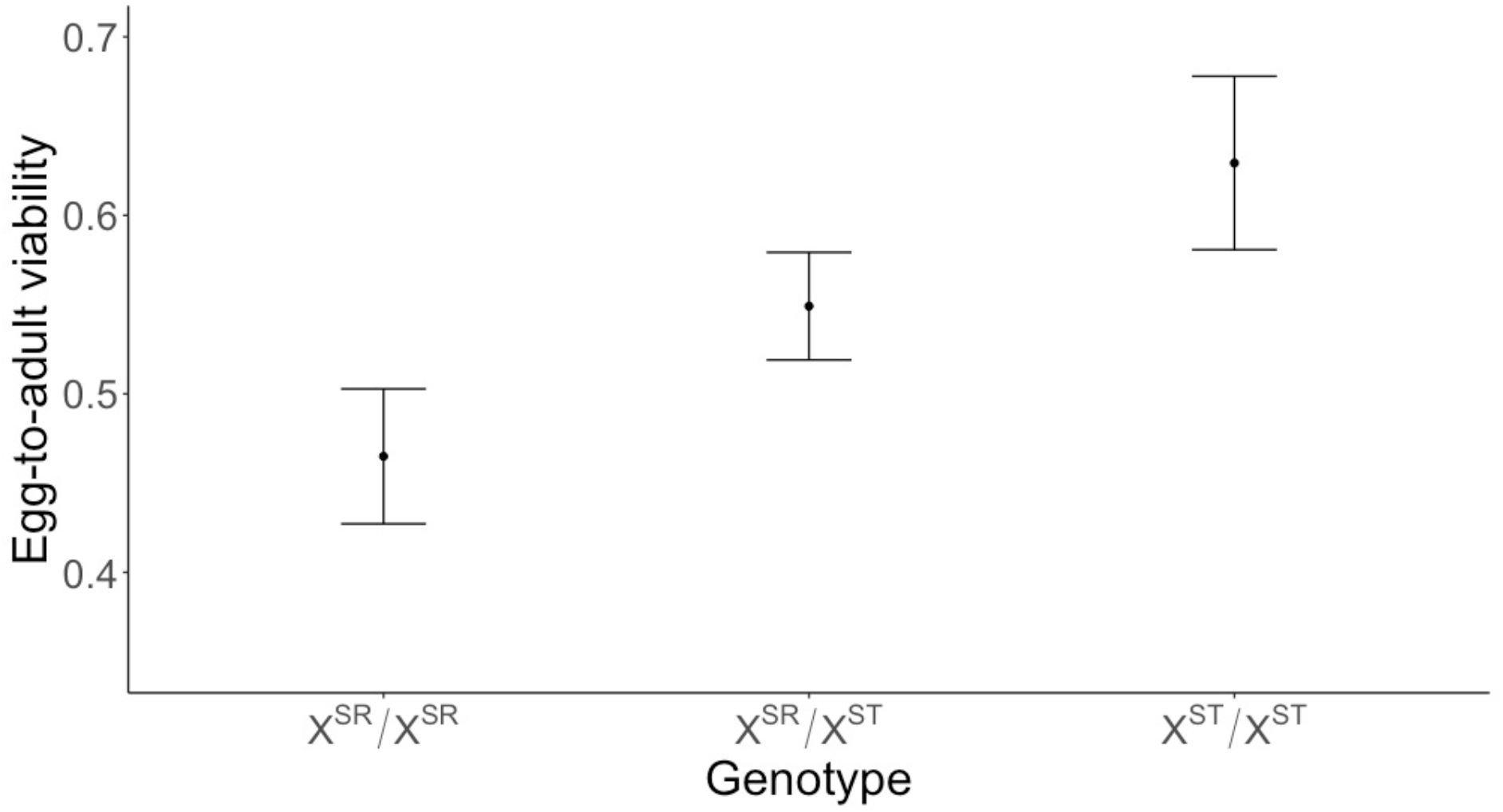
Female X^SR^/X^SR^, X^SR^/X^ST^ and X^ST^/X^ST^ mean ± standard error proportion egg-to-adult viability. Values are determined from the fraction of a given genotype observed in replicate cages.

### Estimating the strength of selection against drive

The posterior probability of each value of the male selection parameter *s*_*m*_ is given in Figure 4. The mode of *s*_*m*_ = 0.214 with a 95% credible interval 0.097 – 0.316 and a 99% credible interval 0.056 – 0.346. The probability of the modal value compared to the null hypothesis of no viability selection against drive males has a Bayes Factor BF_10_ = 321.79.

**Figure 4.**
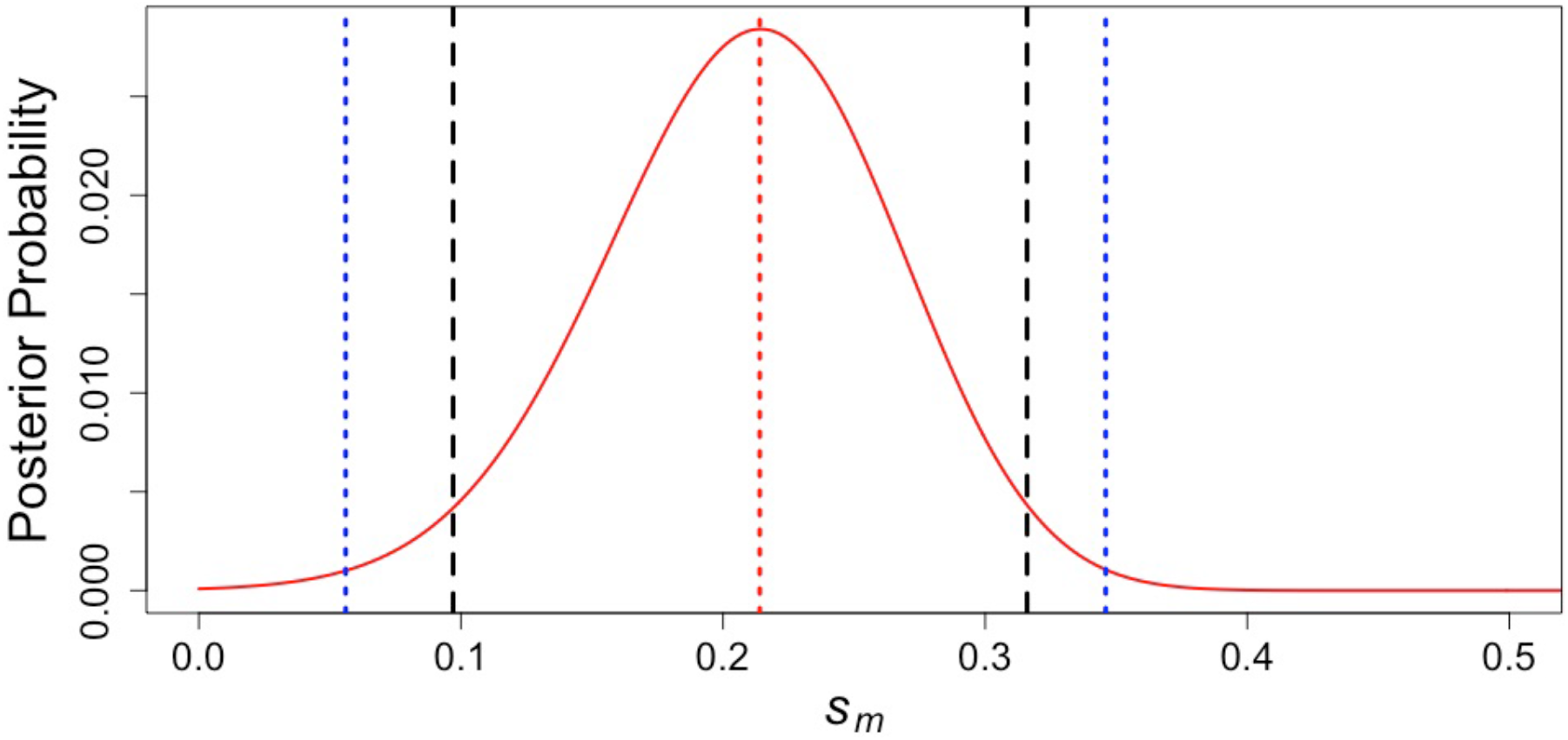
The posterior probability density of the strength of selection against drive in males (*s*_*m*_). The mode is shown as a dotted red line. The dashed black lines indicate the 95% credible interval. The dotted bluelines indicate the 99% credible interval.

The posterior probability of each combination of the female selection parameters *s*_*f*_ and *h* values is shown in Figure 5. The modal values are *s*_*f*_ = 0.242 and *h* = 0.511, with the bivariate 95% and 99% credible interval displayed as a two-dimensional contour (Figure 4). The probability of the modal *s*_*f*_ value compared to the null hypothesis of no viability selection against drive in females has a Bayes Factor BF_10_ = 572.89. The strength of selection against drive in males and females (*s*_*f*_ and *s*_*m*_; setting *h* to its modal value), did not differ between the sexes (|z| = 0.3785 α = 0.01 *P* = 0.7047).

**Figure 5.**
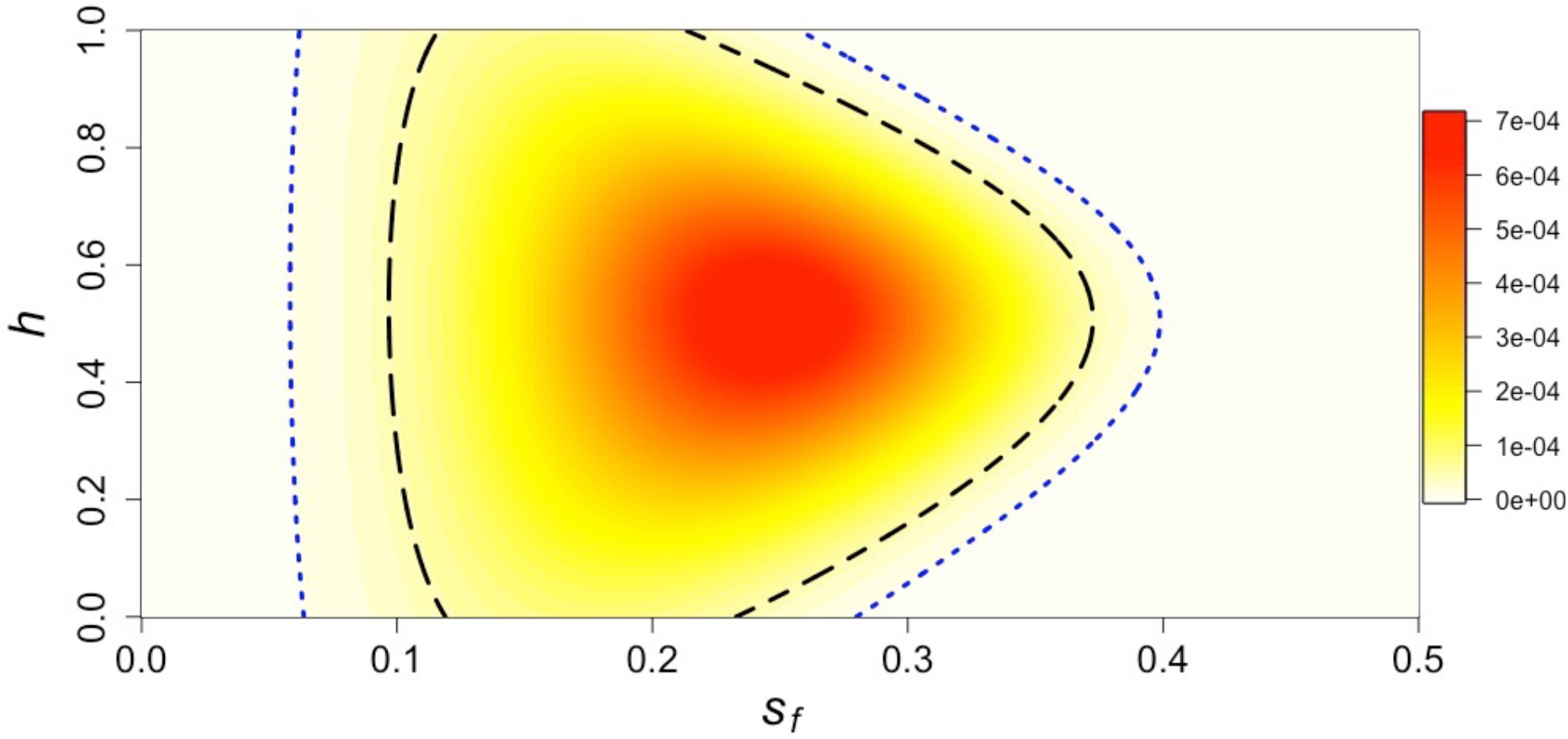
The posterior probability density of the strength of selection against drive in females (*s*_*f*_) and the dominance coefficient (*h*). Colour indicates probability density, with darker colours indicating higher likelihood. The black dashed contour shows the 95% credible interval and the blue dotted line shows the 99% credible interval.

In the pairwise comparison of individual female genotypes there was a difference between the egg-to-adult viability of X^ST^/X^ST^ and X^SR^/X^ST^ females, with a selection coefficient mode = 0.126 with a 95% credible interval = 0.007 – 0.232 and a 99% credible interval = −0.017 – 0.261. A similar difference was observed in the comparison of X^SR^/X^ST^ and X^SR^/X^SR^, with a selection coefficient mode = 0.138 with a 95% credible interval of 0.008 – 0.252 and a 99% credible interval of −0.038 – 0.287.

## Discussion

Due to their two-fold transmission advantage in males, X chromosomes that exhibit *sex-ratio* meiotic drive (X^SR^) potentially can spread to fixation and cause population extinction (Hamilton, 1967; Hatcher et al. 1999). Despite this, several meiotic drive systems exist in broadly stable polymorphisms (Wilkinson et al. 2003; Pinzone and Dyer, 2013; Price et al. 2014). This suggests that there are costs of carrying the X^SR^ chromosome. In the stalk-eyed fly system, the X^SR^ chromosome contains a large inversion (Johns et al. 2005), which is expected to accumulate deleterious mutations as they are less efficiently removed by recombination than those of the X^ST^ chromosome. This mutation load is expected to lead to a decrease in fitness of the X^SR^ chromosome. Here, controlled crosses were used to estimate one component of fitness, egg-to-adult viability, of meiotic drive genotypes. There was a reduction in viability linked to X^SR^ in both males and females. In X^SR^ hemizygous males this was *s*_*m*_ = 21% (Figure 4) and in X^SR^ homozygous females *s*_*f*_ = 24% (Figure 5). The negative effect of X^SR^ in females is largely additive (ℎ ~ 0.5), with heterozygotes being intermediate in viability compared to homozygotes. The estimates of selection (*s*_*m*_ and *s*_*f*_) do not differ between the sexes. This probably reflects a lack of sexual dimorphism in fitness at the larval stage. In *D. melanogaster*, egg-to-adult viability measured for particular genotypes is strongly positively correlated across the sexes, whereas adult reproductive success is typically negatively correlated (Chippendale et al. 2001; Arnqvist and Tuda 2010).

In the experiment, individual males of known genotype, either SR or ST, were crossed with heterozygous females. Eggs were collected and combined in groups of 6 petri dishes each containing 12 eggs. The eggs were visually inspected for signs of development, so as to be able to exclude the possibility that differential fertility of the two paternal genotypes (i.e. SR or ST) affected the subsequent output of adult flies. The combination of eggs from the two crosses were expected to generate all five genotypes in an even ratio, except for heterozygous females which were expected at double the number of the other genotypes. The objective was to standardise competition between genotypes. It is hard to estimate whether this objective was attained, as only surviving adults were genotyped. The observed adult genotype frequencies were compared to infer genotype-specific survival in the egg-to-adult stage. The number of flies genotyped was sufficiently large (*N*_*m*_ = 798, *N*_*f*_ = 1272) to give reasonable assurance of the accuracy of the estimates. Even with this sizeable sample, the bounds on the estimates of *s*_*m*_, *s*_*f*_ and *h* remain large (Figure 4–5) but we can be confident that drive is associated with loss of viability in both sexes. Our results contrast with a prior study of adult lifespan which found no differences in males or females (Wilkinson et al. 2006). The contrasting results may be due to a real difference between larval and adult genotypic effects. But there may have been insufficient power to detect adult genotypic effects as the scale of the adult experiment was one quarter of that used here.

This is the first study showing a reduction in SR viability in stalk-eyed flies. Similar methods have been applied previously in *D. pseudoobscura* (Wallace, 1948; Curtsinger and Feldman, 1980; Beckenbach, 1983). Wallace (1948) observed strong selection against X^SR^ in both sexes. In high density populations, Beckenbach (1983) found a reduction in X^SR^/Y viability but no viability effect on homozygous X^SR^ female viability. In contrast, Curtsinger and Feldman (1980) report stronger selection against homozygous X^SR^ females. Comparisons of these three studies provides strong evidence to suggest that viability selection is density-dependent, as reduction in X^SR^ viability was greatest under high density (Wallace 1948), and a lack of differential viability was observed in another experiment carried out at low density (Beckenbach 1983). In the present study, stalk-eyed fly larvae were cultured under low density and provided with an excess of food. Future work will need to determine whether varying levels of food stress enhance or restrict the deleterious effect of the X^SR^ chromosome.

Strong viability selection against the X^SR^ chromosome, as found here under laboratory conditions, will play a key role in determining the equilibrium level of the SR polymorphism in the wild. There are several other factors that could be involved in determining SR frequency, such as suppressors, polyandry and various forms of sexual behaviour which we discuss further here. First, in *D. simulans*, SR commonly co-occurrs with suppressors which restrict the transmission advantage (Merçot et al. 1995; Kingan et al 2010). Although early work on the stalk-eyed fly drive system suggested that there were suppressors (Wilkinson et al. 1998), this has not been sustained by further work, either on the autosomes or Y chromosomes (Paczolt et al. 2017). Second, polyandry may evolve to limit the spread of SR (Price et al. 2008). Polyandry is the norm in *T. dalmanni* (Baker et al. 2001; Wilkinson et al. 2003), and there is evidence that SR male sperm does less well under sperm competition (Wilkinson et al. 2006) and may suffer from interactions with non-sperm ejaculate components produced by standard males (though this has only been shown in the related species *T. whitei*, Wilkinson and Fry 2001). But it has not been shown whether elevated polyandry occurs in populations of *T. dalmanni* with higher frequencies of SR or in stalk-eyed fly species that harbour drive (compared to those that lack drive).

Third, it has long been suggested that mate choice may play a role in determining the frequency of drive (Coopersmith and Lenington, 1990). This may be important in stalk-eyed flies as they are canonical examples of sexual selection driven by mate choice (Burkhardt and de la Motte 1983; 1985). In *T. dalmanni*, drive males are expected to attract fewer females as they have reduced eyespan, and hence mate less often (Wilkinson et al. 1998; Cotton et al. 2014). However, there is as yet no evidence in stalk-eyed flies that the strength of female mate preference has been enhanced in populations subject to drive. Nor has there been investigation of whether females that carry SR show alterations in their mating behaviour. A related consideration is male mate preference (Bonduriansky 2001) which has been shown to be an important behavioural adaptation in *T. dalmanni* favouring male matings with fecund females (Cotton et al. 2015). A recent study reported that SR had no direct effect on male mate choice (Finnegan et al. 2019). However, the strength of male mate preference positively covaries with male eyespan. As drive males have smaller eyespan (Cotton et al. 2014), we expect they will be less discriminating in their mate choice (Finnegan et al. 2019).

Finally, measurements of sperm number per mating report that SR males deliver as many sperm as ST males, and a single mating with a SR male results in the same female fertility as a mating with a ST male (Meade et al. 2018). Whether this pattern carries over to situations where a male can mate with multiple females is less clear. One experiment showed no difference between SR and ST males (Meade et al. 2019), whereas another experiment found lower fertility in SR males (Wilkinson et al. 2003) when multiple females were allowed to mate freely with a single male for a day. The cause of this difference is unclear, but drive males have been shown to have lower mating rates compared to standard males (Meade et al. 2019), and this could conceivably have contributed to lower fertility in females mated to SR males. As mentioned previously, P2 experiments indicate that SR males are poor sperm competitors with ST males which must arise from reasons other than numerical sperm transfer from the male (Wilkinson et al. 2006).

The number of different factors makes it difficult to predict the equilibrium frequency of drive in the wild and whether these factors are sufficient to explain the observed frequency of ~20% (Wilkinson et al. 2003; Paczolt et al. 2017). Many could act as stabilizing forces which restrict the spread of drive in a frequency-dependent manner. Future work should aim to examine these factors, in combination with the intensity of egg-to-adult viability selection measured here, in a modelling framework in order to predict the evolutionary outcomes. This can then be related to better estimation of parameters across local populations of *T. dalmanni* in which SR frequency is known to be highly variable (Cotton et al. 2015) along with experimental evaluation of interactions between the various male and female enhance or constrain the various selective forces.

## Acknowledgements

SRF is supported by a London NERC DTP PhD Studentship (NE/L002485/1). AP is supported by Engineering and Physical Sciences Research Council grants (EP/F500351/1, EP/I017909/1), KF and AP by NERC grants (NE/G00563X/1, NE/R010579/1). The authors thank Rebecca Finlay for her help in maintaining the laboratory stocks, Dr Lara Meade for advice on experimental design and Prof Mark Thomas for help with the statistical analysis.

## Data accessibility statement

Data will be made available at the Dryad Digital Repository

## Author contributions

The research project was conceived by SRF, NJW, KF and AP. The experiment was carried out by SRF and NJW, with genotyping by SRF, DK and MFC. The data was analysed by SRF, NJW and AP, and the paper written by SRF and AP.

